# CUDA RNAfold

**DOI:** 10.1101/298885

**Authors:** W. B. Langdon, Ronny Lorenz

## Abstract

We add CUDA GPU C program code to RNAfold to enable both it to be run on nVidia gaming graphics hardware and so that many thousands of RNA secondary structures can be computed in parallel. RNAfold predicts the folding pattern for RNA molecules by using O(*n*^3^) dynamic programming matrices to minimise the free energy of treating them as a sequence of bases. We benchmark RNAfold on RNA STRAND and artificial sequences of upto 30 000 bases on two GPUs and a GPGPU Tesla. The speed up is variable but up to 14 times.

## 1 Introduction

The central dogma of Biology [Crick, 1970] says that the fundamental information for all forms of life is transcribed from DNA into messenger RNA, which in turn is translated into protein. Like DNA, RNA is a long chain polymer principally composed of 4 bases (G,U,A, and C) also like DNA the four bases can form relatively weak temporary bonds with their complementary base. (C pairs with G, and A with U) allowing an RNA molecule to fold up on itself. This is known as its secondary structure. Much of the chemistry of biomolecules is determined by their three dimensional shape. In the case of RNA, unlike chemically more diverse molecules like proteins, the secondary structure can help when trying to infer or explain RNA molecule’s biological impact [Fallmann *et al.*, 2017]. Figure 1 shows the secondary structure of a 20 base RNA molecule. (The right hand side is a rendering of one of its three dimensional shapes [Chen *et al.*, 2006, Fig. 1].)

**Figure 1:**
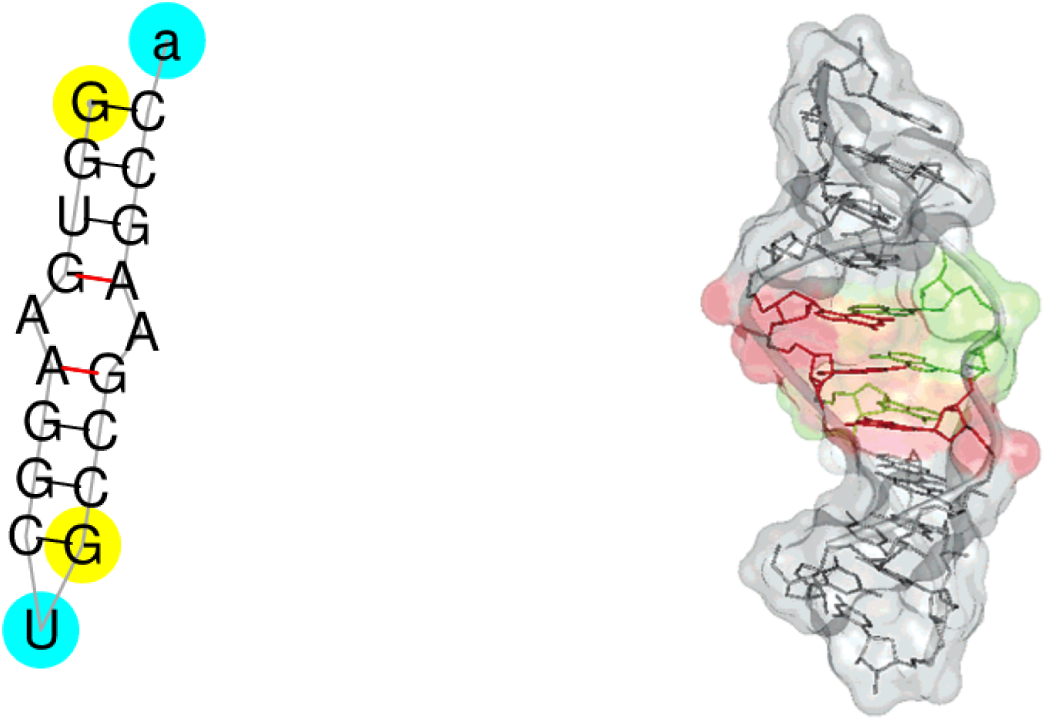
Folding pattern (secondary structure) for RNA molecule PDB 00996 (sequence GGUGAAG-GCUGCCGAAGCCa). Left: secondary structure as recorded by RNA STRAND v2.0 [*Andronescu et al., 2008*]. Red lines indicate non-standard RNA base-to-base binding, which RNAfold assumes does not happen. Right: one of true three dimensional structures [Chen *et al.*, 2006].

RNA Secondary structures are typically computed under the assumption of isolation, i.e. fixed ion concentration and no bound ligands, and thermal equilibrium. Thermal equilibrium in particular then dictates that the probability of a particular structure is proportional to the Boltzmann weight of its free energy, and so the structure with minimum free energy (MFE) is the most probable one. To compute such properties, efficient dynamic programming algorithms based on a Nearest Neighbour (NN) energy model exist. They decompose RNA structures into loops of unpaired nucleotides delimited by so-called canonical base pair, each associated with a free energy term derived from experimental data and mathematical models based on polymer theory. However, to allow for efficient recursive decomposition and due to the scarcity of energy parameters, certain structural configurations are by definition excluded from secondary structures. This includes knots and pseudo knots, though they might, albeit rarely, appear in natural RNAs [Andronescu *et al.*, 2008, Table 1]. The program RNAfold, from the ViennaRNA package [Lorenz *et al.*, 2011], is the state of the art C code which predicts how an arbitrary RNA molecule (represent as a string of G,U,A, and C) will fold up. It is widely used. For example, it is a key component of the eteRNA citizen science project [Lee *et al.*, 2014].

**Table 1:**
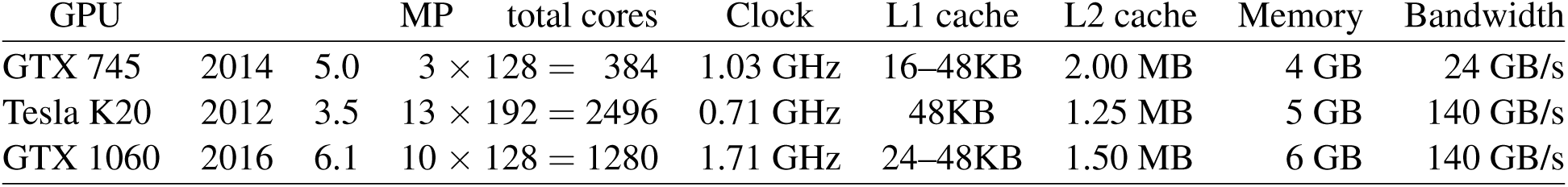
GPU Hardware. The year each GPU was announced by nVidia is in column 2. Third column its compute capability level. Each GPU chip contains 3, 10 or 13 identical independent multiprocessors (MP, column 4). Each MP contains 128 or 192 stream processors. (Total given in column 6). Clock speed is given in column 7. The next two columns give sizes of the L1 and L2 caches. These on chip cache sizes are followed by (off chip) on board memory and, in the last column, the maximum measured data rate (ECC enabled) between the GPU and its on board memory.

RNAfold is based on using dynamic programming to calculate every feasible folding pattern and select the one with minimum energy. (By looking only at “feasible” patterns the number of possibilities is constrained and dynamic programming general scales as O(*n*^3^) (where *n* is the length of the RNA molecule).

Except in short RNA, the bulk of RNAfold’s computational demands come from its use of dynamic programming. It calculates the energy to be minimised using symmetric matrices, where the interaction energy between the *i*^th^ and *j*^th^ bases is held in matrix element *ij*. Since this is the same as *ji* only half each matrix need be stored. The main memory requirements of RNAfold come from needing at least three upper-triangular matrices per sequence, two of which relate to computations that will be delegated to the GPU. As modern GPUs have multiple gigabyte of on board memory, the CUDAGPU version of RNAfold can process *< G/*4*n*^2^ sequences in parallel. (Where *G* is the GPU memory size in bytes and *n* is the length of the RNA sequences. 4 converts size in bytes to size in integers.) E.g. a4GBGTX745 can process upto about 26 000 RNA molecules of size 200 nucleotides in parallel^1^.

The next section will describe the initial analysis which lead to the discovery of how to break RNAfold’s highly nested dynamic programming code into substantial parts that could be run in parallel and the decision to create GPU code for the largest of these: modular decomposition (Section 3.1) and int loop (Section 3.2). The changes to the existing host code are sketched in Section 3.3. The new code’s performance on three diverse GPUs on real RNA (actually the whole of RNA STRAND) and both shorter and longer RNA sequences are given in Section 4. Section 6 summarises and discusses how the CUDA versions of RNAfold and pknotsRG compare. Some earlier work is described in Section 5 Whilst Appendix A describes the implementation in more detail and gives hints on CUDA development is general.

## 2 Planning: Profiling with GNU gcov

The ViennaRNA package (release 2.3.0) was downloaded and installed on our 3.6 GHz desktop. Much of the profiling, development and benchmarking was done on natural RNA whose structure is known using 4661 RNA molecules from RNA STRAND v2.0 [Andronescu *et al.*, 2008]. Figure 2 gives one of the larger examples.

Except where noted, we run RNAfold with its default options. The GNU GCOV profiling tool confirms that for larger RNA molecules almost all the computational effort is taken by the dynamic programming in program source file mfe.c. It confirmed the importance of modular decomposition loop, which we had previously used genetic improvement to optimise with Intel SSE 128 bit instructions [Langdon and Lorenz, 2017], and the fill arrays code in general.

**Figure 2:**
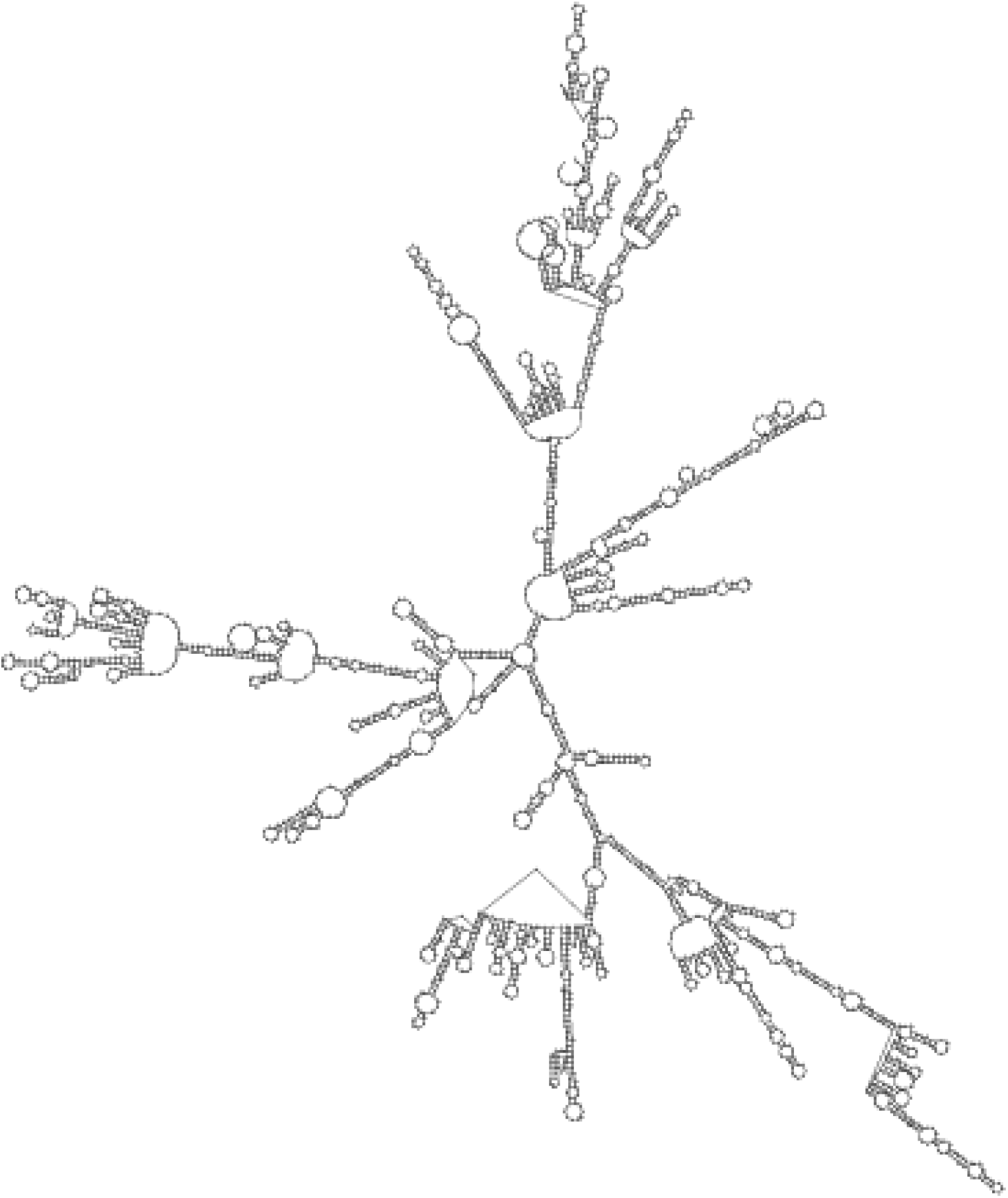
Predicted secondary structure of 2913 long RNA molecule from Spinach 23S Ribosome (RNA STRAND V2.0 sequence CRW 01456).

## 3 Code changes

Using the output of GCOV for guidance the existing release 2.3.0 C code was split into two new include files fill arrays.c and fill arrays loops.c The existing sequential code relies on data from one operation being available for the next, making it hard to exploit the natural parallelism in Dynamic Programming. The new code was progressively re-written to split the nested for loops in to related lines of code calculating intermediate energy values which could be isolated and moved out of the main inner *j* loop but pass data into it via arrays rather than scalars. GCOV showed that many of these calculations were not computationally demanding and so were (for the time being) left on the host CPU rather than being calculated on the GPU.

This left two main calculations at the inner most (*j*) loop, each of which is now done by a CUDA kernel on the GPU. That is, each time the kernel is called by the host for a particular *i* value, it processes all of the corresponding *j* iterations in parallel. Typically this is about (*n − i*) *×* nsequences operations.

To minimise data transfers, the GPU maintains a copy of two corresponding “triangular” half matrices: my fML and my_c. They each occupy (*n* + 1)(*n* + 2)*/*2 int per sequence (where each sequence is*n* bases long). I.e. the space required on the GPU is about 4*n*^2^*×* the number of sequences bytes in total.

To enable the remaining non-GPU operations to be run, the host and GPU copies of both my fML and my c must be kept in step. This is done once per *i* loop by copying the changes across the PCIe bus linking the CPU and the GPU. A number of small CUDA kernels are used to do this and to initialise data structures.

### *3.1* modular decomposition

The modular decomposition kernel essentially replicates the modular decomposition mfe.c code, see Figure 3. It is run once per *i* loop, replicating a complete *j* loop, which itself invokes a still deeper loop in the original host code. The deepest loop adds two values from my fML, one from a column and one at the same height from the diagonal. It then steps one element down takes the sum and then takes the minimum of the two sums, until it reaches the base, by which point it has found the minimum for that column which is inserted into a vector of new energy values. It then moves on to do the same on the next column.

**Figure 3:**
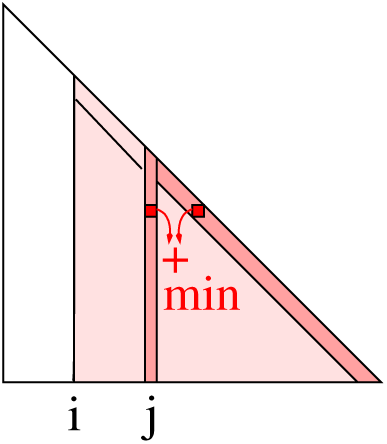
Ignoring end effects, to get the *j*^th^ output for step *i*, modular decomposition adds the intermediate energy value stored on the matrix diagonal to that at the same height in the *j*^th^ column and then takes the minimum of all such sums. The calculation is repeated for every *j* value bigger than *i*, thus sweeping the whole of the lightly shaded data. The GPU version does this in parallel. Only the updates are passed back to the host CPU. The next time modular decomposition is used, *i* will be one smaller (i.e. have moved one place to the left) and the contents of the triangular matrix will have been updated.

The only slight complication, is that if either value to be added has not been calculated (i.e. it has the special value INF) then that pair is skipped. If all values are skipped the output is set to INF. The simplicity of the calculation, fundamentally means the modular decomposition parallel calculation on the GPU is limited by reading and writing the data within the GPU (see last column in Table 1) rather than the GPU’s calculation speed.

On a GPU it is possible to do these calculations in parallel. modular decomposition uses BLOCK SIZE (BLOCK SIZE is currently tuned to be 64) threads to do 64 sums and minimisations in parallel. When all are done it uses a shared memory reduction to yield the smallest of (up to) 64 minimums. NB in principle all threads are needed by the reduction stage, even when the value they have calculated is INF. One block of threads do all the calculations needed for a single output value. This avoids atomic operations (which have a high latency overhead). All the other outputs are calculated in parallel by other blocks.

Each time modular decomposition kernel is launched, it will read all of the triangular matrix above the *i*^th^ column. In general the triangular matrix is far too big to be held in rapid access cache memory. For cache performance reasons (as on the CPU), it turns out to be better to use another kernel, fmli kernel, to copy the most active part my fML (i.e. the diagonal) into a separate small array before launching modular decomposition kernel. The data in the small array is used many times but may be small enough that the GPU can keep it all in L2/L1 cache.

The kernel header is written to use the __restrict__C++ keyword on all pointer arguments. This potentially allows the compiler to make additional optimisations. In the case of computer level 3.5 GPUs (i.e. the Tesla K20, see Table 1) the nVidia compiler may access read only data (i.e. arguments marked with the const keyword) via a small texture cache.

The appendix (Section A.2) contains a reasonable quantitative simple model which uses arguments of bandwidth and cache sizes to predicts the performance of modular decomposition kernel.

### 3.2 int_loop

Like modular decomposition.cu, int loop.cu is a new file which holds all the GPU code to process internal loops. It too works principally on one large triangular matrix, my_c. my_c is the same size as modular decomposition.cu’s my fML (i.e. *n*(*n* + 1)*/*2). The algorithmic code to calculate intermediate energy values is based on interior loops.h function IntLoop X (release ViennaRNA-2.3.0). On the GPU, IntLoop_X is called by __device__function Energy, which itself is called from CUDA kernel int_loop_kernel. Like modular decomposition kernel, ignoring end effects, int loop kernel is launched for each column *i* of my_c, starting at the largest value (near *n*), control returns to the host CPU, and the next time modular decomposition kernel is launched *i* is one smaller. NB. as in Figure 3, as *i* gets progressively smaller, the length of the *i*^th^ column, and so the volume of calculations, increases. Ignoring edge effects, each time int loop kernel is launched it calculates new values for all the *j* elements in column *i* of my_c. Unlike modular_decomposition_kernel, int_loop_kernel essentially works with a local part of its main input my_c.Ignoring edge effects, for each value of *j* it performs calculations on a square region 31 by 31 (actually 0 … MAXLOOP) of my_c, see Figure 4. Notice, depending on hard constraints, adjacent values of *j* will use overlapping squares regions of my_c. Thus when run to calculate column *i*, int loop kernel only needs access to columns (*i*… *i* + 31) of my_c. This locality should mean up to about *i* cache lines (128 bytes each) are occupied by my_c when int_loop_kernel is active.

**Figure 4:**
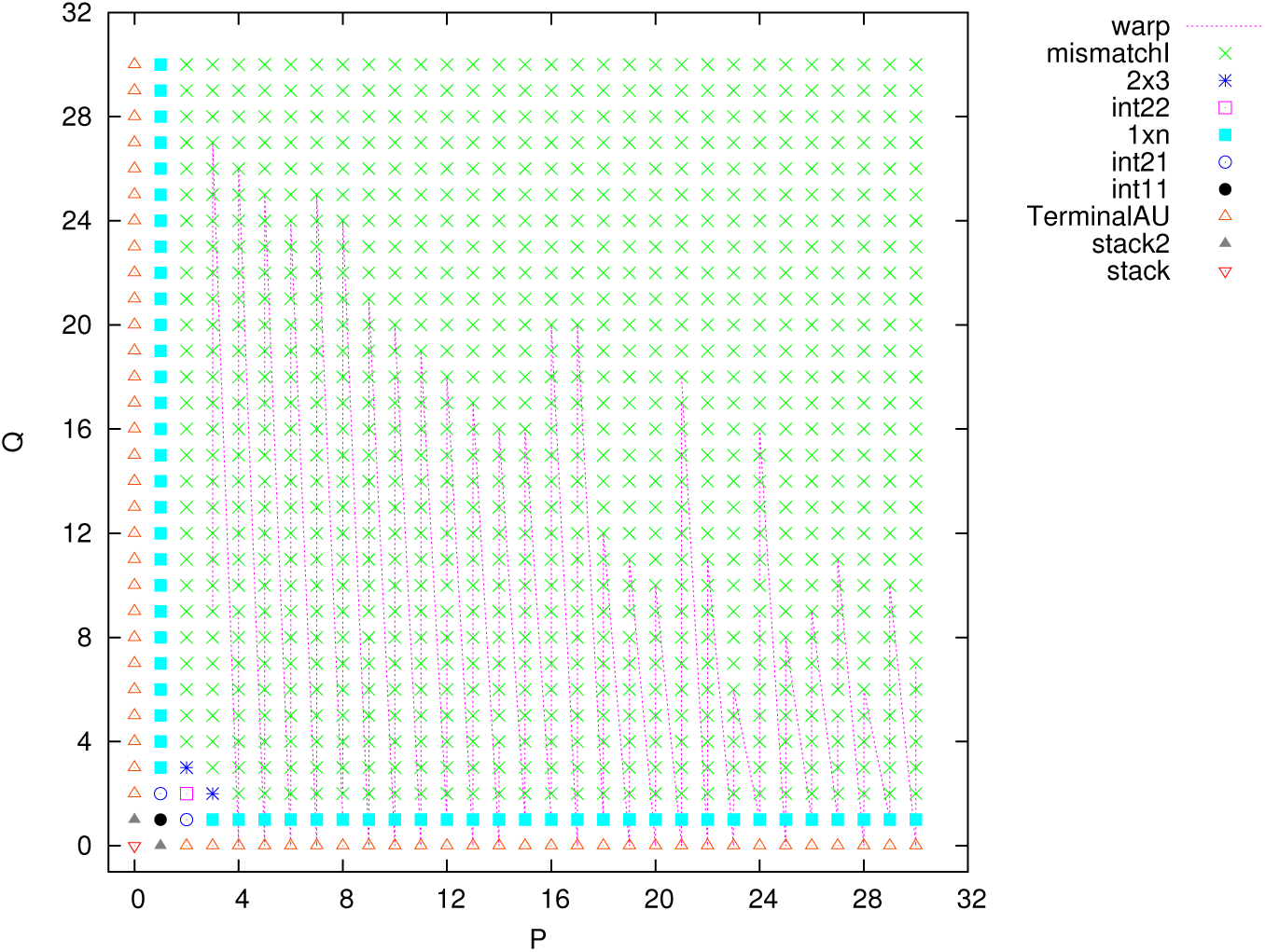
Example calculation of internal loop intermediate energies by IntLoop X. The different crosses correspond to the ten different routes through IntLoop X (both int21 routes shown with blue ⊙). Due to the hard constraint VRNA CONSTRAINT CONTEXT INT LOOP ENC, of the 31^2^ = 992possible values only 478 are calculated. The new device routine setpq picks these out (dotted lines) so only 15 warps are needed (15 32 = 480) rather than 31. In this example in all cases the warp of 32 threads diverges because more than one path is included.

Like modular_decomposition_kernel, int_loop_kernel need perform its calculation only for array elements if my_c is not INF, but also each calculation must be skipped unless the hard constraint is satisfied. In principle for each output cell up to 31^2^ intermediate calculations must be made and added to the corresponding element in myc and then the output cell is set to the minimum of all of them.int_loop_kernel does these calculations in parallel. In principle we could use up to 31^2^ different threads to do as much work as possible in parallel, however initial experiments suggested that using 32 threads gives the fastest calculation. Figure 4 shows an example where the 32 threads are spread across my_c. In the example each thread performs 14 or 15 calculations and updates its own local instance of the minimum. Like modular_decomposition_kernel, a parallel reduction using shared memory is used to find the minimum of these 32 minima and then a single thread is used to write the global minimum to the output. Again using a single block of threads per output allows the calculation to be done in parallel with minimal synchronisation and without the use of expensive atomic operations.

Figure 4 shows that each *i*,*j* pair can take one of ten routes through the code in IntLoop X. Typicallyeach route does little calculation but must read data from a different source. In Figure 4 each *i*,*j* pair is labelled with either the name of the loop or the major data structure that must be read. These data are RNAfold parameters [Langdon *et al.*, 2018]. Only those used by intLoop X are stored on the GPU, occupying 201728 bytes of global memory. We anticipate that, on modern GPUs, most blocks of code running int_loop_kernel will find them already in L2 cache.

Typically due to hard constraints only about 30% of the 31^2^ = 961 values need be calculated. On the host the hard constraints are pre-calculated and stored as a bit mask in char array. Since only one bit of the mask is used on the GPU, to reduce memory bandwidth we extract this bit on the host and store it in a bit packed array d hccc on the GPU (access to d_hccc is via__device__function Hc). We experiments with different layouts of the d_hccc bit array, but surprisingly found performance to be relatively impervious to row v. column order.

The paths through IntLoop_X are given by geometry and so can be precalculated. We introduced __device__ function setpq to ensure only internal loop energies meeting the hard constraints are calculated. Using 32 threads in parallel setpq finds all the set bits in d hccc for the 31 by 31 square region and so causes IntLoop X to be called just for them. Each group of 32 threads, known as a warp, is linked by a dotted line in Figure 4.

setpq minimises the number of threads with little work to do but proved complex to code. As Figure 4 shows typically each warp passes through more than one branch in IntLoop X. This means the threads in the warp diverge and so do not exploit the full power of the GPU. Since each branch will read different data, this divergence is perhaps more severe than divergence just due to requiring different code to be executed. However other mappings of *i*,*j* (including symmetric herring bone patterns) were tried, and despite the complexity of coding them, they gave no noticeable performance improvement. Another approach was to code a unique kernel corresponding to each branch in IntLoop X. Initial results were not promising due to the cost of starting kernels which might have little work to do (e.g. int11). However perhaps this approach warrants further investigation.

If we are right and the data read by int_loop_kernel is in L2 cache, we struggle to explain why its performance, even on modest GPUs, is so low.

### 3.3 RNAfold.c *and* mfe.c

The CUDA code will by default choose the fastest GPU available. The routine new routine choose gpu(argc,argv) allows the user to select another GPU via the device command line option. It also starts the CUDA code. (Perhaps this CUDA warm up code could be in a separate pthread.)

In order to be able to process multiple sequences in parallel the existing while loop which read sequences was modified slightly so that instead of calling vrna mfe(vc, structure) immediately the data created for each sequences are saved in arrays. Only when the last sequence has been read in and process is a new routine par mfe(nfiles,VC,Structure,EN) called.

For ease of implementation the intermediate arrays are of fixed size MAXCUDAPAR (65535). Since these arrays only contains pointers, there is actually little overhead on creating the arrays bigger than the actual number of sequences (nfiles) to be processed. These five arrays are given upper case names corresponding to the original scalar variable, whose values they store.

To aid quick implementation, the code to interactively analyse RNA sequences has simply been commented out. Similarly, as yet, there is no attempt to support pf.

New routine par mfe (in mfe.c) essential initialises the GPU code, and emulates vrna mfe as if it had been called once per sequences (i.e. nfiles times). However where vrna mfe would have called fill arrays the new code calls new routine par_fill_arrays. par_fill_arrays is in new include file fill arrays.c^2^ and is based on fill arrays, however it also includes additional sanity checks. These do not appear to be especially onerous, however, once the code is debugged, for efficiency, the repeated calls to sanity(VC[0],VC[H]) could perhaps be omitted.

Although the initialisation code in par fill arrays is fairly standard, the main i and j loops have been re-organised and re-ordered to enable much of inner j loop to be do in parallel. In our case, this is done by parallel CUDA kernels on the GPU but the restructuring could potentially be applied to other parallel architectures. Also there are a few parts of the original j loop that remain on the host CPU but which could be ported to the GPU.

At present the key data structures are maintained instep on both the host and the GPU for each interaction of the outer (i) loop. If the remainder of the j loop processing were moved to the GPU, potentially the small overhead in keeping the host and GPU instep could be reduced. However at present this does not look like it would give a big payback.

Some (unused?) non-default options in the original fill arrays code have not been supported as yet. Instead their code is often commented out and/or protected by assert statements.

## 4 Results

The characteristics of the three systems used are given in Tables 1 and 2. Figure 5 shows the CUDA code can be considerably faster than the original code but there can be a large start up overhead. The time taken to start the GTX 1060 (180 milliseconds) and the Tesla K20 (413 milliseconds) may be due to some CUDA misconfiguration. (The corresponding time is 145 milliseconds for the GTX 745.) ^3^

**Figure 5:**
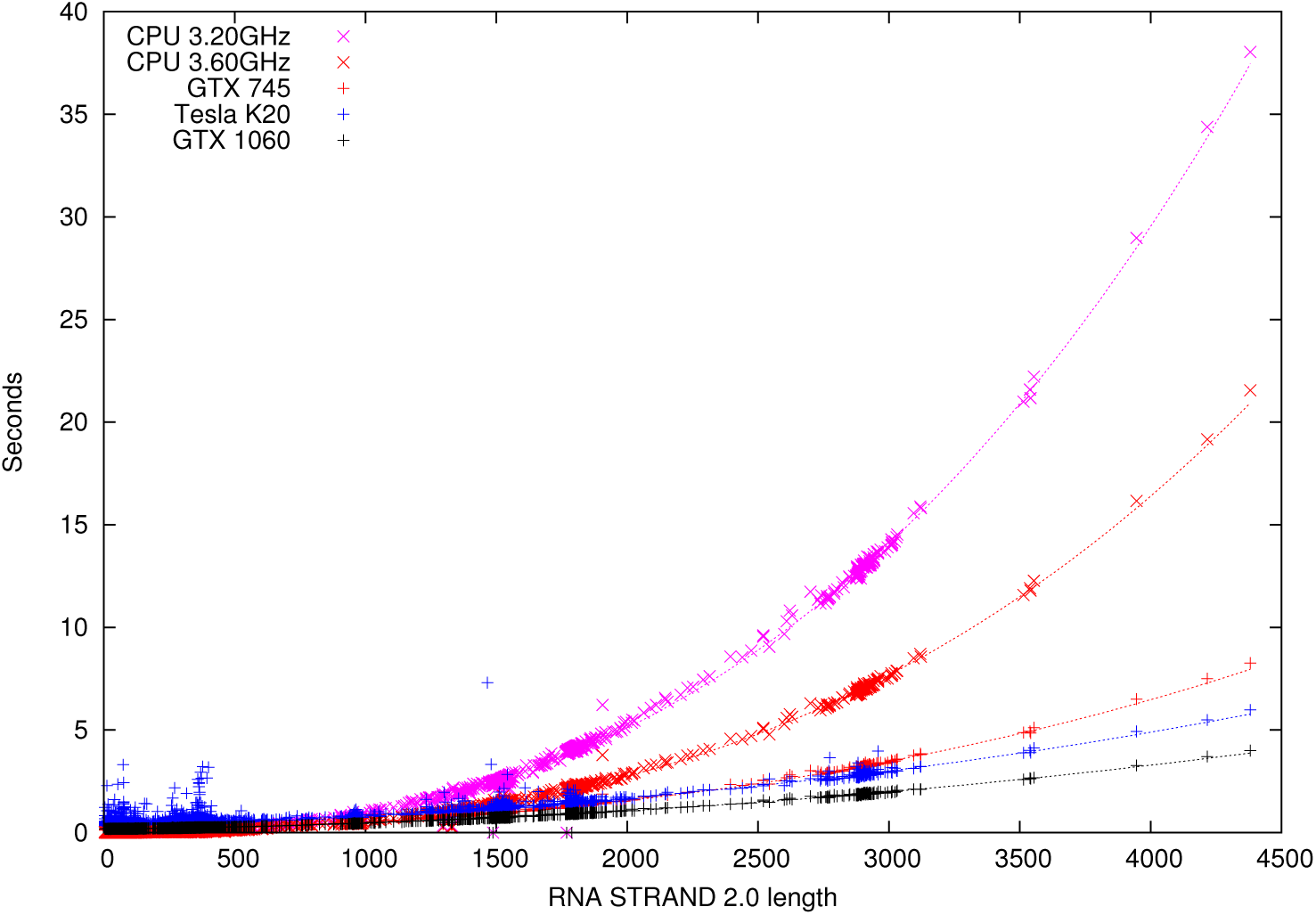
Time taken to process real RNA. 4662 RNA molecules from RNA STRAND V2.0. In all cases our CUDA version of RNAfold produces exactly the same predicted secondary structure (or fails in the same way as RNAfold release 2.3.0). Dotted lines show best cubic fit.

**Table 2:** CPUs. The GTX 745 (top row) and GTX 1060 (last row) are each housed in desktop computers, whilst the Tesla K20 is in a computer server.

| Type | Cores | Clock | Memory |
| --- | --- | --- | --- |
| Desktop | 8 | 3.60GHz | 31 GB |
| Server | 8 | 2.67GHz | 12 GB |
| Desktop | 4 | 3.20GHz | 3 GB |

Since RNAfold is used, e.g. by the eteRNA project, to predict unknown RNA structures, we also bench-marked it on many unknown random RNA sequences. Figure 6 shows, except for run times ≪1 second, that the computation time for random RNA sequences is the same as for natural ones, which is expected.

**Figure 6:**
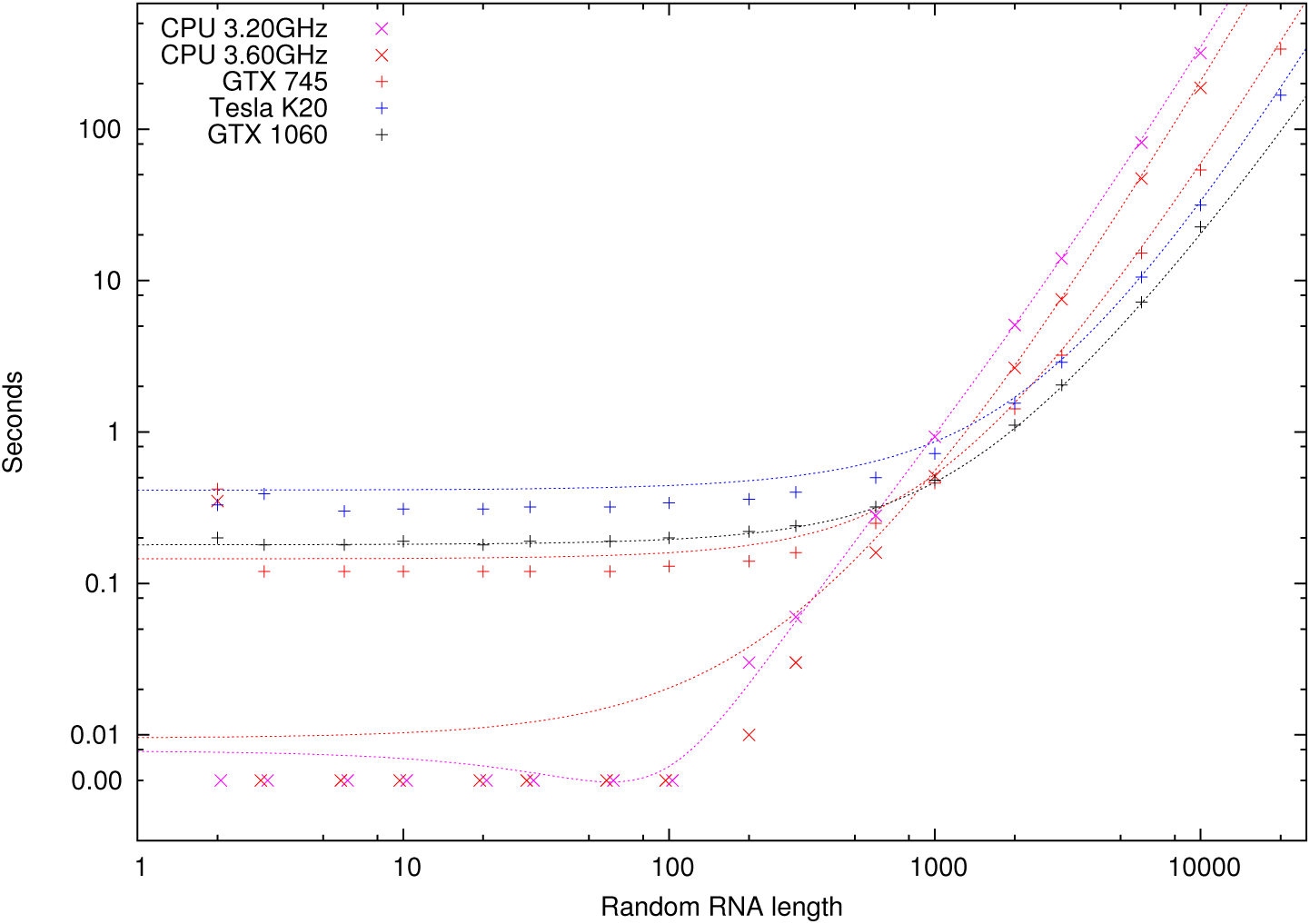
Time taken to predict RNAfolding (note log scales). Dotted lines show are best fit for RNA STRAND V2.0, i.e. same lines as in Figure 5. Data below one second are proportionately more noisy.

As mentioned on page 2, the CUDA version can process many files in parallel. Figure 7 expands Figure 6 by plotting the total run time to predict the secondary structure of many sequences in parallel. (Figure 8 shows the same data but expressed as a speed up over RNAfold, release 2.3.0.) Particularly with shorter sequences, running many RNA molecules in parallel gives an improvement over running them one at a time. (See Figure 9.To reduce clutter Figure 9 shows data only for one GPU.)

**Figure 7:**
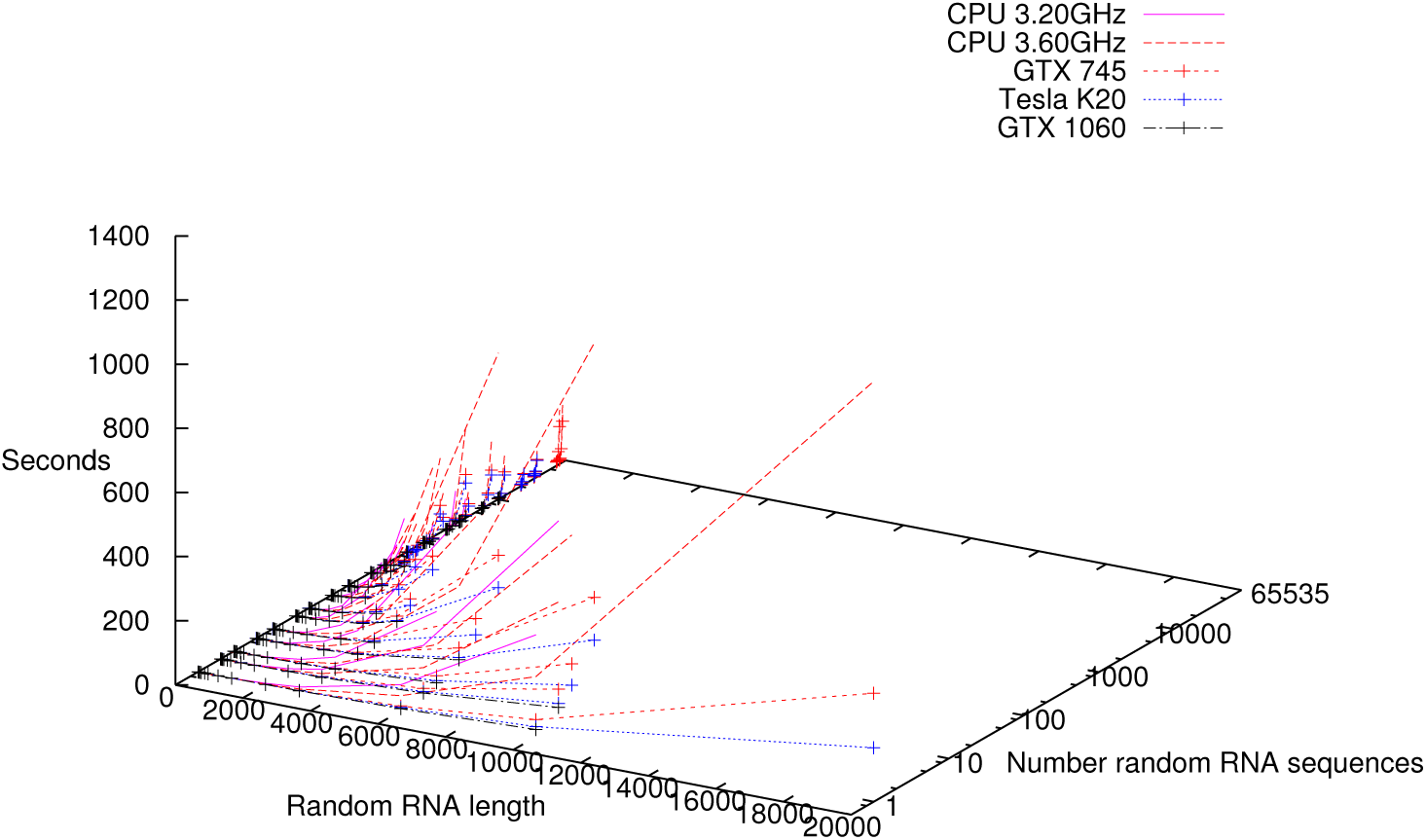
Time taken to predict RNA folding (note number of sequences processed in parallel plotted on a log scale).

**Figure 8:**
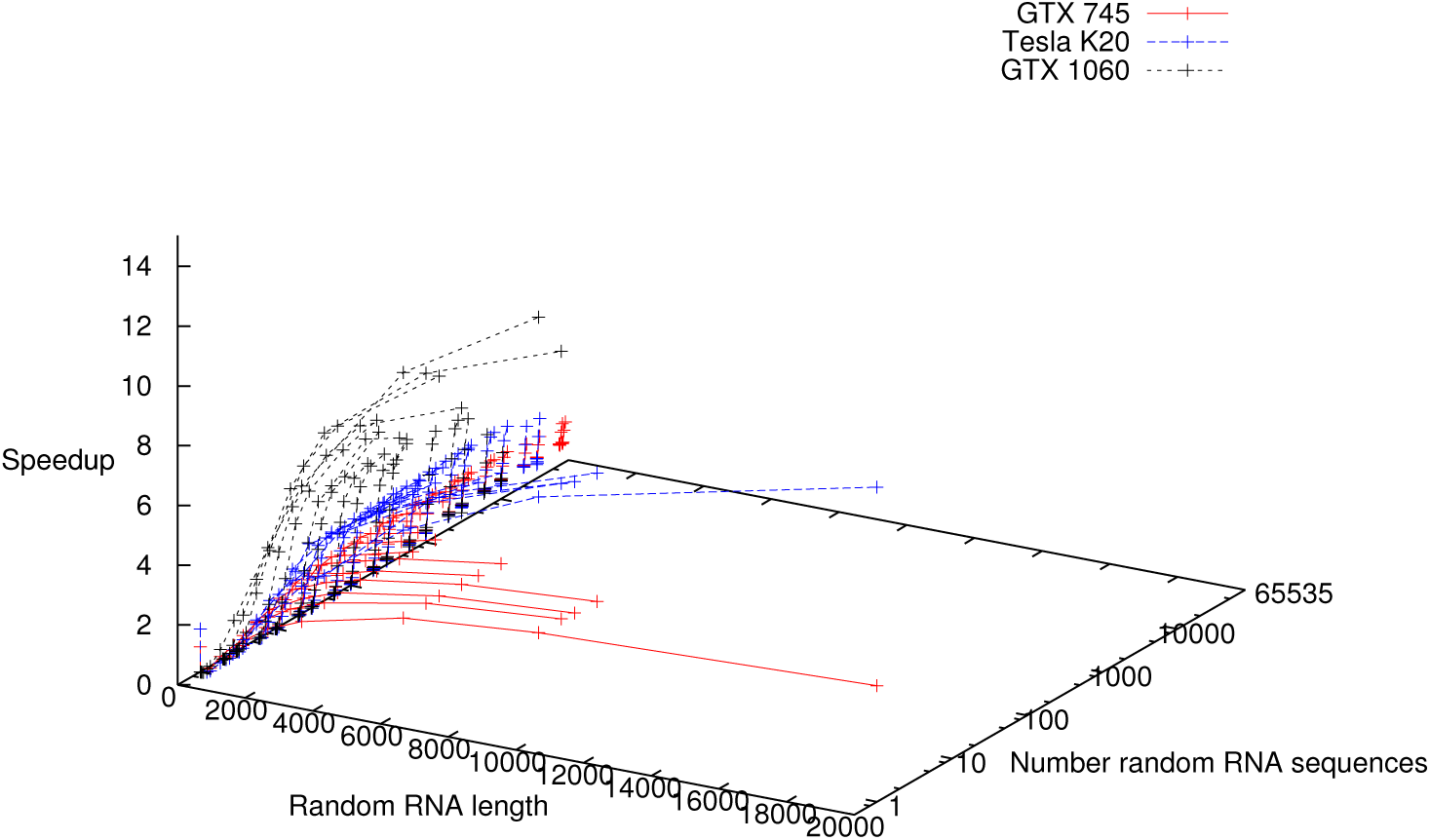
Speed up GPU v. one core on host (note number of sequences processed in parallel plotted on a log scale). Same data as Figure 7. Due to LTO problems (Section A.3), CPU time for K20 server estimated from CPU time for desktop hosting GTX 745 using ratio of their CPU clocks (Table 2).

**Figure 9:**
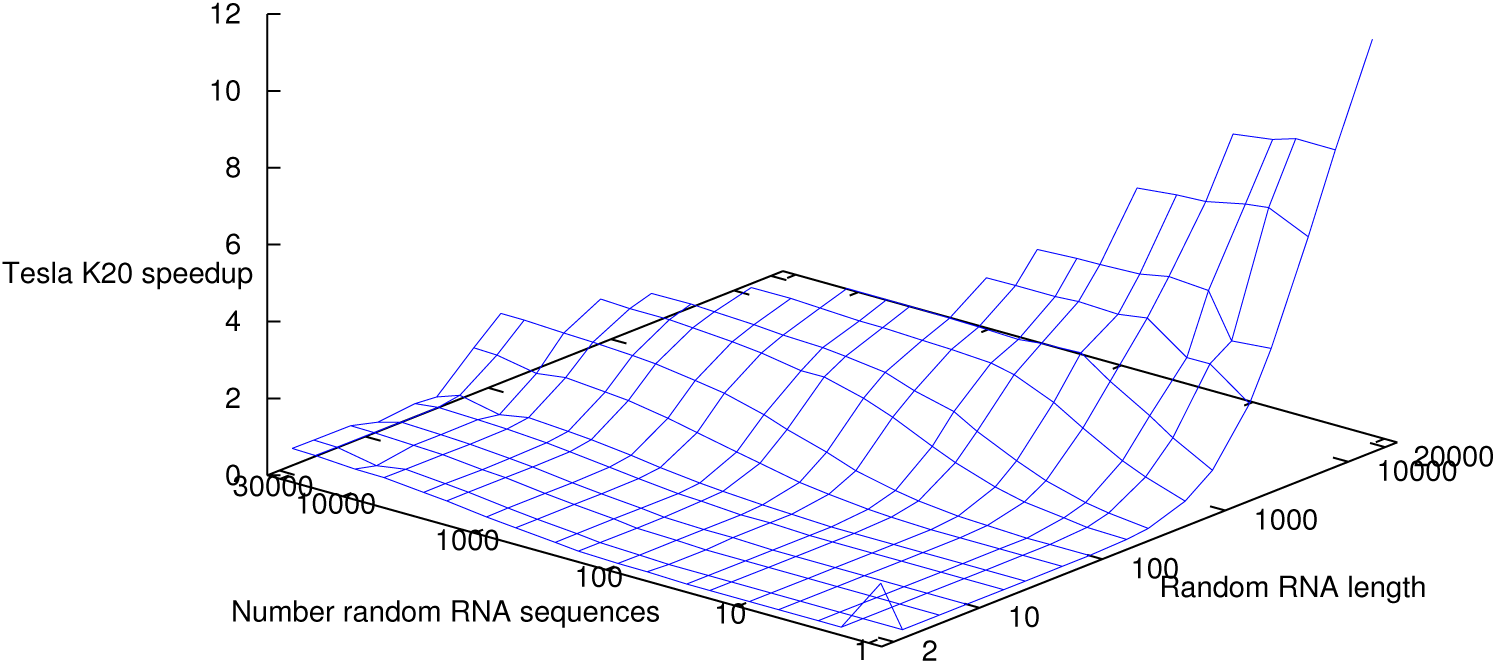
Speed up Tesla K20 v. one core on host (note log scales). Data as Figure 8.

## 5 Prior work

We have benchmarked RNAfold on the whole of RNA STRAND as this is effectively all the RNA molecules whose true structure is known. As these tend to be small (median length 294), like [Stern and Mathews, 2013], we have also used random RNA sequences to benchmark the computer programs. As Figure 6 shows, these fit the average trend of RNA STRAND well. In some cases, earlier work has omitted various molecules in RNA STRAND as they have unusual folding behaviour or pseudo knots. Another approach is to benchmark computer programs on longer real RNA molecules (whose structure need not be completely known) such as those associated with HIV [Hofacker *et al.*, 1996]. 20 years ago [Hofacker *et al*., 1996, p24] considered the Ebola virus (≈ 20 000 bases) might one day be within the range of of their super computer, we have demonstrated today 20 000 bases is well within the reach of domestic consumer electronics.

Unlike [Stern and Mathews, 2013], our GPU version produces identical answers to the original code. For speed [Stern and Mathews, 2013] converted their O(*n*^3^) code from double to single precision to run it on their CUDA GPU. They take care to show the loss of accuracy is small. Since we’vefocused onlyon minimum free energy prediction, almost all corresponding calculations are made as standard 32 bit integers. Like [Stern and Mathews, 2013, p6] we determine the best thread block size empirically. The longest sequence [Stern and Mathews, 2013, Tab. 1] report is 9709 (GenBank AF324493). We took the first 9709 bases from AF324493.1 (complete sequence HIV-1 vector pNL4-3)^4^ [Stern and Mathews, 2013, p^7^] report their Tesla M2050 (448 cores) took 27 minutes to predict base pair probabilities for it, whereas on our slowest GPU, it takes less than a minute to predict the best folding. (However bear in mind, although both algorithms are O(*n*^3^), [Stern and Mathews, 2013]’s requires more multiplications than are needed to just predict RNA folding.)

Like us [Lei *et al.*, 2012] based their implementation on CUDA and on code taken from the ViennaRNA package. It appears that their implementation is based on taking advantage of the GPU’s shared memory, which is very fast but in short supply (≤48KB). Therefore it appears that their system limits the size of RNA molecules it can process. At any rate they present results only for four lengths (68, 120, 154 and 221 bases, see * in Figure 10). Also they only present mean performance calculated when thousands of random RNA sequences of the same size are processed together. As is common, we have presented speed ups comparing a GPU v. a single CPU core. Although performance on Intel CPUs typically scale sublinearly with number of cores, comparison with a single core is a widely used convenient baseline. Performance on other cores and clock speeds can be guestimated from it. However [Lei *et al.*, 2012] present most of their GPU data as ratios compared to a complete 2.4 GHz personal computer with four cores, and so the speed ups they present are even more impressive. Although they have a sophisticated system to balance the load between the 4 CPU cores and their nVidia GTX 280 GPU it appears to have provided at best only a 16% advantage on their benchmarks over the GPU itself. ^5^

**Figure 10:**
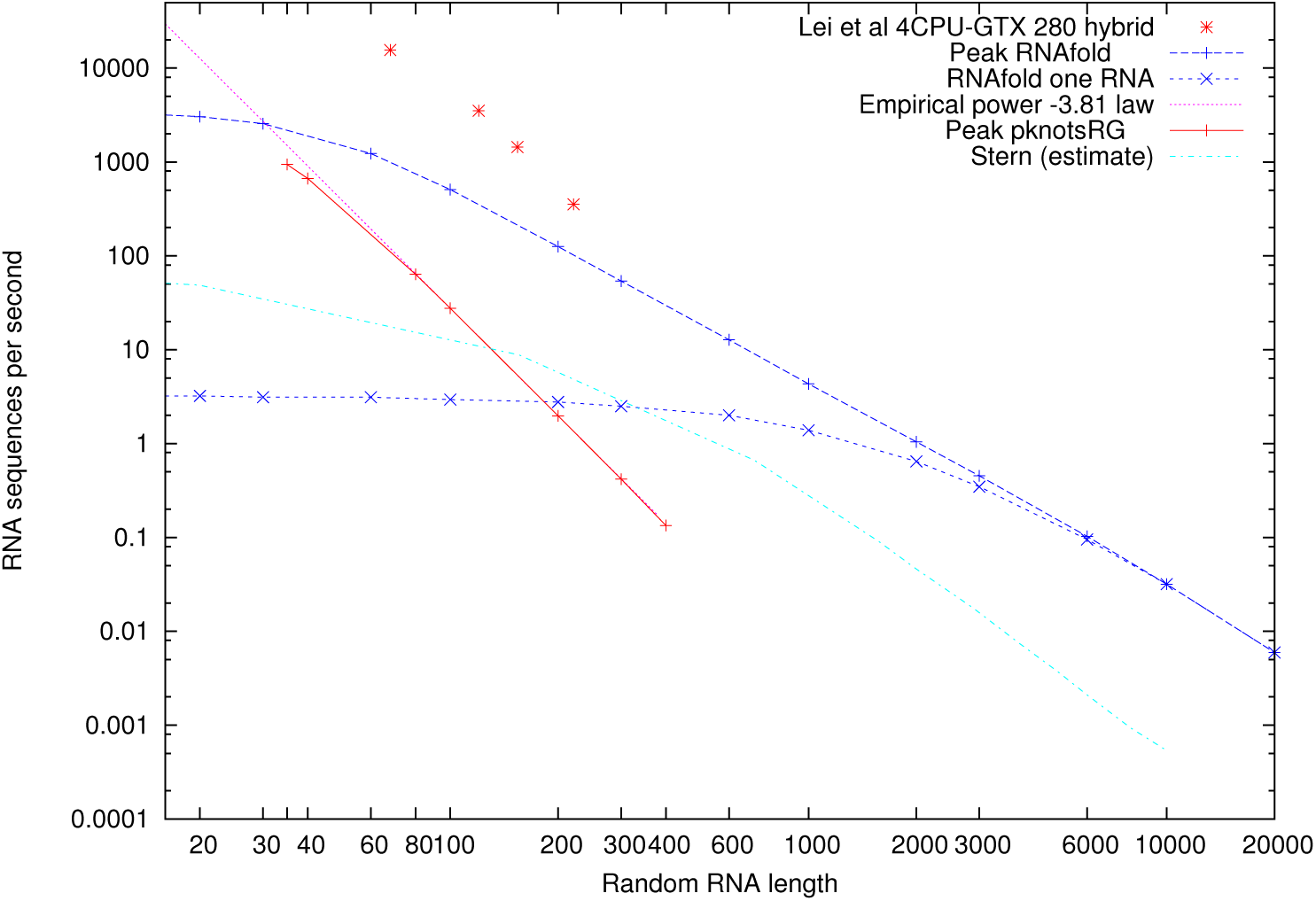
CUDA versions of pknotsRG (solid line), RNAfold (dashed lines), [Lei *et al.*, 2012]’s CPU-GPU hybrid (*) and [Stern and Mathews, 2013] (light blue line). pknotsRG K20 data from [Langdon and Harman, 2015a, Fig. 4]. Notice pknotsRG is solving a different, albeit related, problem. It includes recursive pseudoknots prediction, for which the asymptotic complexity is O(*n*^4^), which compares well with the observed scaling of *n^−^*^3.81^. While RNAfold omits all pseudoknots and has complexity O(*n*^3^). CUDA RNAfold data as Tesla K20 in Figure 7. For comparison we also present average speeds from [Lei *et al.*, 2012, Tab. 4] for 20 000 random RNA sequences. Data are only available for the four lengths (68, 120, 154, 221) plotted. Plot for [Stern and Mathews, 2013] estimated from [Stern and Mathews, 2013, Fig. 2] using WebPlotDigitizer 4.1

### 5.1 Evolving RNAfold

Grow and graft genetic programming (GGGP) [Harman *et al.*, 2014; Jia *et al.*, 2015; Kocsis and Swan, 2018; Langdon *et al.*, 2015; Langdon and Harman, 2015a; Langdon, 2015a; Langdon *et al.*, 2017] has built on existing genetic improvement (GI) work [Langdon, 2015b; Petke *et al.*,]. Genetic Improvement has been used to improve the performance of existing software, e.g. by reducing runtime [Langdon and Harman, 2015b], energy [Bruce *et al.*, 2015] or memory consumption [Wu *et al.*, 2015], but (apart from software transplanting [Marginean *et al.*, 2015] and automatic bug fixing [Le Goues *et al.*, 2012]) it usually tries not to change programs’ output. GGGP builds on Genetic Improvement which is very much hands off (i.e. let the computer solve the problem) by admitting more human assistance. pknotsRG, like RNAfold, was an open source C program for predicting the secondary structure of RNA. However, unlike RNAfold, it came with an existing CUDA version. It was this version which using GGGP we obtained enormous speed ups [Langdon and Harman, 2015a]. The original CUDA kernel was automatically generated but, like the C version of pknotsRG, could only process one structure at a time. Our version obtained huge speed ups on quite small RNA molecules by processing many of them in parallel (see solid line in Figure 10). Unfortunately pknotsRG is buggy and no longer supported and so we switched to the state of the art code, RNAfold, which is being actively maintained.

We applied GGGP to RNAfold [Langdon and Lorenz, 2017]. Using traditional manual methods to identify performance critical components, recoding them using Intel’s SSE vector instructions and then using computational evolution to further improve the new code. Unfortunately, evolution only found small increments on the human code and so, so far, only the manually written code has been adopted. However it has been included into the standard ViennaRNA package since 2.3.5 (14 April 2017)^6^. It is also being used by the eteRNA development team internally [Das, 2017].

We have also applied GGGP to improve the accuracy of RNA fold’s predictions [Langdon *et al.*, 2018]. Notice there we allowed (actually encouraged, required) evolution to change the output of RNAfold, whereas here the CUDA version produces identical answers.

## 6 Conclusions

We have demonstrated the new version of RNAfold gives identical answers to the official released code from which it was derived and yet can give speed ups of up to fourteen fold on modern consumer video gaming GPUs.

This work was inspired by the enormous speed up we obtained for the CUDA version of pknotsRG [Langdon and Harman, 2015a]. The great speed up came from the ability to predict the structure of many RNA molecules simultaneously. Our new CUDA version of RNAfold can also make many thousands of structure predictions in parallel but,even for short RNA sequences, it gives more modest returns. This disparity may be because for short sequences, RNAfold’s serial (O(*n*)) preprocessing is a large fraction of the total run time. In contrast pknotsRG did little serial preprocessing. Instead it contained a single CUDA kernel, which did almost all the dynamic programming work. Whereas for our CUDA version of RNAfold, with a single long sequence, the dynamic programming (O(*n*^3^)) processing is already saturating the GPU.

## Acknowledgements

I am grateful for the assistance of Bobby R. Bruce, Neil Daeche, UCL, Derek Jones, Knowledge Software, UK, Rhiju Das, Jukka Suomela, shaklee3, callum.burns, BulatZiganshin, njuffa and txbob (of nVidia).

The K20 Tesla was donated by nVidia.

## A CUDA Implementation Lessons

Unfortunately CUDA programming remains far from straightforward. In the hope that lessons learnt may be useful, perhaps to other projects, we give some details of the RNAfold CUDA implementation process.

### A.1 Processing Multiple RNA sequences in Parallel

The serial part of RNAfold (RNAfold.c and mfe.c) was tweaked to allow multiple sequences to be run in parallel giving a substantial speed up compared to running one sequence at a time (For example, in the case of the Tesla K20, cf. dashed and dotted lines in Figure 10). Unlike our earlier work with pknotsRG [Langdon and Harman, 2015a] the extra sequences are not tightly integrated and simply result in additional thread blocks being run in parallel. Data such as the RNAfold parameters are common and so are shared by multiple sequences and need only be loaded once. We see a big performance boost, probably due to the start up overhead being shared by many sequences.

### A.2 Optimal Performance

The major runtime components remain:

- On the host CPU the time taken to read and sequentially process RNA sequences.
- Initialising the GPU (start up overhead described on page 7).
- int_loop_kernel takes about two thirds of the GPU run time.
- modular_decomposition_kernel takes approximately one third of the GPU run time.

In the case of modular_decomposition_kernel we can estimate the best performance we should get based on the size of the data to be read [Giles, 2017, Lecture 2, p. 9]. (See Figure 3.) Each block of modular_decomposition_kernel uses a parallel for loop to read int energy values from a small linear array and add them to similar data from a large triangular array and writes the minimum such sum to a linear int array of energy values. The small array is re-used by every block but each block reads a different part of the large array. To estimate the best possible performance we assume the small array is always in cache and that the kernel’s runtime is dominated by the time to read the rest of the data. Assuming *n >* 9, each block reads 1 to *n* — 9 int and modular_decomposition_kernel uses *n*9 blocks,so data read = 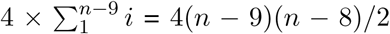bytes.For example for *n* = 3000, on our GTX 745 (cf. Table 1) reading the large array should take at least 720 microseconds, which compares very well with an observed runtime of 799 microseconds. Figure 11 shows as expected run time is O(*n*^2^). NB. remember that the modular decomposition code will be run *n* times, giving the expected O(*n*^3^) overall run time.

**Figure 11:**
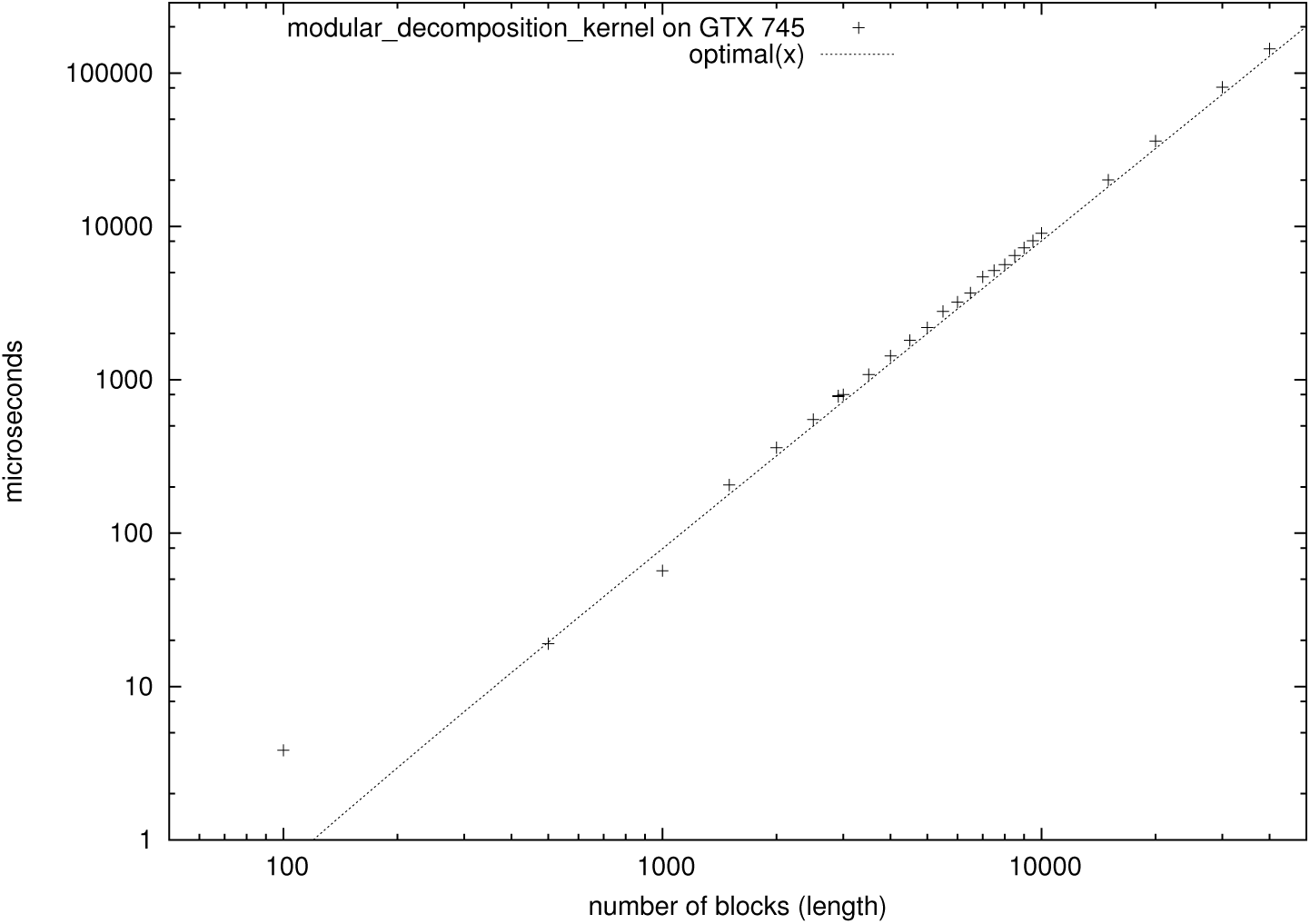
Observed and predicted best possible run time for modular_decomposition_kernel on GTX 745.

We did not manage to get a satisfactory explanation for the performance of the int_loop_kernel.

### A.3 No GNU Link Time Optimisation (LTO)

At present the nVidia compiler is not compatible with the GNU GCC’s link time optimisation^7^ In some cases it seems sufficient to link with default switches but in other cases it seems necessary to explicitly turn it off (e.g. using -Xcompiler -fno-lto). The ViennaRNA package makes extensive use of various -f -lto switches during compilation and so disabling LTO might have an important impact on the performance of serial code remaining on the host. A potential work round would be to extract the CUDA code to a separate link time image. The auxiliary CUDA image could still share data with the non-CUDA image, so the runtime overhead of separate linking might not be excessive.

### A.4 Code optimisations

#### A.4.1 Floating point

There is very little floating point calculation in RNAfold. We did however take the precaution of following the usual C programming convention to ensure all floating point calculations are make in single precision and floating point constants are similarly stored as float rather than double.

#### A.4.2 *No* -use fast math

As expected there was little impact on using the nVidia compiler option -use_fast_math and so for simplicity it was omitted.

### A.5 CUDA tools

#### A.5.1 source code printf, assert

printf, assert can be planted into the GPU source code and work much like the corresponding C tools.

With printf you must be aware that it is run in parallel and can easily generate vast amounts of data. Firstly there may be far too much output to be useful but also it is easy to overflow the GPU’s print output buffer. Hence if a large volume of debug output is generated, and an output is missing you should consider if it might have been lost as well that it might not have been generated in the first place. Secondly the GPU does not ensure that data written in parallel appear on stdout in the same order as they are generated. Hence it is usually best to generate complete lines of output (i.e. up to a \n) even if the line might at first appear unwieldy.

As with host code, the GPU assert can be a useful debugging tool. However it is common practise for efficiency (as sometimes with host CPU code) to use the compile time switch -DNDEBUG to disable it. Thus disabling all assert based sanity checks. It may be worth using conventional if …exit code with one time host based sanity checks to avoid this.

#### A.5.2 CUDA Memcheck

A lot of GPU code uses pointers to access data structures. cuda-memcheck is a good run time tool for checking pointers indicate valid parts of GPU arrays and data structures. It can be applied to a program after it has been compiled and linked but does impose a large run time overhead.

Since GPU warps of 2^5^ threads are automatically synchronised a common coding pattern is for efficiency during the last five steps of a shared memory reduction not to limit the bounds read by each thread. This leads to half the threads reading outside the bounds of the allocated shared memory. Since their output will be ignored, this has no impact on the output but cuda-memcheck may spot this and issue an error message.

#### A.5.3 nVidia Profiler

Another useful run time tool is nvprof. It too can be applied after a program is run and linked. It gives limited summary data about the number of times CUDA operations and CUDA kernels are run and the time they took. Unlike cuda-memcheck it only imposes a very modest run time overhead^8^.

#### A.5.4 nVidia Visual profiler

Unlike nvprof nvvp is designed for interactive use. It seems to impose only modest overhead. nvvp will usually need to run the program multiple times, for usability it is important to find *short representative* program run before starting to use nvprof. nvprof is applied to a program after it has been compiled and linked. Depending on versions, it may be able to exploit information planted by the compiler for debugging purposes at compile time but this it is not essential to compile with these switches.

Whereas nvprof is simple to use and gives a single output, nvprof has many options and multiple displays. It is easy to get very lost in the vast number of data it generates and lose track of the message they are trying to convey. Perhaps the most useful display is the time line which shows when major events took place, how long they took and the gaps between them.

### A.6 CUDA Help

All the CUDA documentation is online. Additionally nVidia support several online user fora. Perhaps the most directly useful is the forum on “CUDA Programming and Performance”.

nVidia continue to improve their tools. For example, nvvp is now considerably more stable than the first released version. Also, on the GTX 745, we found a 10% performance improvement on migrating from CUDA 7.0 to 9.1.

1 To simplify coding, CUDA RNAfold uses thread grids. Typically this imposes its own limit of 65535 sequences. However this is only relevant to GPUs with large memories or to short RNA sequences and could be relaxed if needbe.

2 New include file fill arrays.c similarly includes new file fill arrays loop.c

3 The original start-up times for both the GTX 1060 and TeslaK20 were much worse (i.e. 0.88 and 3.00 seconds) due to lack of CUDA persistence mode. The improved times are given by using, as a workaround, --persistence=1 on an image run in the background simply to keep the CUDA driver in state where it is ready for immediate use. See discussion at https://devtalk.nvidia.com/default/topic/1030608/cuda-programming-and-performance/why-2-9-seconds-to-start-tesla-k20/ The problem has been raised with nVidia, bug ID: 2081242.

4 We took version 1 from GenBank as it was current when [Stern and Mathews, 2013] was published.

5 [Lei *et al.*, 2012]’s port to Intel SSE instructions appears to have been disappointing and in most cases did not yield a speed up. In contrast our earlier SSE port ([Langdon and Lorenz, 2017] see Section 5.1) gives a modest but consistent speed up of 30% and has been distributed with the ViennaRNA package since release 2.3.5 (14 April 2017).

6 To use the SSE code, ViennaRNA must be configured with ./configure --enable-sse when it is compiled. https://www.tbi.univie.ac.at/RNA/documentation.html

7 https://devtalk.nvidia.com/member/2249473/ Posted 01/11/2018 06:42 PM. The problem has been raised with nVidia, bug ID: 2081246.

8 https://devtalk.nvidia.com/default/topic/1029890/reduce-overhead-of-launching-a-new-thread-block/#5239796 Posted 02/11/2018 04:06 PM

